# Global invasion patterns and dynamics of disease vector mosquitoes

**DOI:** 10.1101/2024.08.06.606755

**Authors:** Rebecca Pabst, Carla A. Sousa, Franz Essl, Adrián García-Rodríguez, Daijun Liu, Bernd Lenzner, Anna Schertler, José Luís Zêzere, César Capinha

## Abstract

Mosquitoes are regarded as the most dangerous creatures on earth, spreading deadly pathogens through their bites. Human activities are driving range expansions of many mosquito species by unintentionally introducing them beyond their native ranges. Despite the often dire consequences for human health, a global picture of the introduction trends and the resulting range expansions of mosquitoes is missing. Here, we describe the global invasion patterns of mosquitoes that are vectors of human diseases and analyze the drivers shaping them. In addition, we provide the dataset compiled for these analyses which represent the most up-to-date standardized information on first records for this taxonomic group at a regional level. Our findings reveal that a total of 45 mosquito species have hitherto been introduced into regions outside their native range worldwide, representing 24% of those known to transmit human pathogens in the wild (i.e., outside experimental conditions), with 27 species successfully established. There has been a steep increase in introductions of emerging non-native mosquito species since the mid-20^th^ century when 28 species (62% of all introduced species) were recorded for the first time. In just the last two decades, 12 new species have been identified. The geography of introductions largely mirrors global trade and transportation flows. Initially, most introduced species were native to Africa, but over time, Asian species have become more dominant. North America, Australia and Europe have consistently been the primary recipients. Our results provide a foundation for addressing the increasing threat of non-native vector mosquitoes globally, emphasizing the need for international cooperation and comprehensive control measures to mitigate their impact on public health.

## Introduction

Human activities often lead to the unintentional introduction of species beyond their native ranges^1,2^. While this phenomenon has existed for millennia, the rate of species introductions, particularly intercontinentally, has accelerated over time^3^. Especially the European exploration and colonialism in the late 15^th^ century contributed to the large-scale redistribution of biota^4,5^. This rate has further increased in recent centuries due to the expansion of transportation networks and the globalization of trade and travel^6–9^. Introduced species can profoundly affect local biodiversity ^10^, human economic activities^11–13^ and pose major threats to human health^14–16^.

The global distribution and spread dynamics of non-native species have been extensively studied in recent years^2,3,17^. However, somewhat surprisingly, there remains a significant knowledge gap regarding the global invasion patterns and dynamics of disease vector mosquitoes. Non-native disease vector mosquitoes are currently found worldwide, altering disease transmission patterns and leading to serious consequences for public health^18,19^. Epidemics of diseases such as Zika and dengue are now common in tropical and subtropical regions due to the high abundances of competent non-native vectors^18,20,21^. In temperate regions, these vectors are also causing a rise in the cases of autochthonous transmission (i.e., acquired infection where transmission occurs between individuals through the local mosquito population)^22,23^. Additionally, due to their substantial impact on public health systems, mosquitoes are among the most costly non-native species worldwide^11^. Reported costs to human societies of mosquitoes of the Genus *Aedes* and the diseases caused by the arboviruses they transmit rises to an aggregated value of 94.7 US$ billion (2022) accumulated over the period 1975–2020^13^.

The history of non-native mosquito introductions is both long and complex. Due to their limited active dispersal ability, mosquitoes primarily spread regionally through human activities (e.g., trade and transportation)^24–26^. The first recorded invasions date back to early intercontinental maritime and trading activities as early as 1495, involving species such as *Aedes aegypti* and those from the *Culex pipiens* complex^24^, which have been primarily dispersed through water-carrying containers on ships^27,28^. The subsequent intensification of global trade, development of the shipping industry and expansion of aerial and terrestrial transportation networks, coupled with a growing human population, has further exacerbated the opportunities for mosquito spread^24^. Adult mosquitoes, their larvae, and their eggs can now hitch rides on planes^29^, cargo ships^25^, and terrestrial vehicles^30^. Additionally, urbanization is also creating artificial breeding sites, such as standing water in discarded containers, facilitating the establishment of some newly arrived species^31^. One of the most notable examples is *Ae. albopictus,* an Asian-native species that has spread globally through the trade of commodities such as used tires and lucky bamboo plants, particularly during the 20^th^ century^26^. Very recently, several emerging non-native mosquito species have been reported worldwide. For instance, *Ae. japonicus*, native to East Asia, has spread to North America and Europe within the last few decades^32^, while *Anopheles stephensi*, native to South Asia, has recently been detected at the Horn of Africa, posing a new threat for malaria transmission in urban settings^33,34^.

The current global invasion patterns and dynamics of mosquito vectors of human diseases remain largely unknown, due to incomplete knowledge about natural and anthropogenic dispersal mechanisms. The spread dynamics of this group have been primarily assessed for single species and only for those causing the most severe impacts^26,35,36^. Studies that have examined the invasion of mosquito species, have focused on mapping species-specific distributions^e.g., 26^, are geographically restricted^e.g., 37^ or are outdated^e.g., 18,24^. Understanding the dynamics of introductions of all non-native disease vector mosquitoes, as well as the patterns and drivers of their spread and establishment, is crucial for developing effective surveillance control measures to prevent future invasions.

To fill these gaps and track the introduction of non-native mosquito species that are vectors of human diseases, and to provide a basis for informed predictions about future invasions and their potential impact on human health, we have compiled a database of their regional first records worldwide through a literature search. We then used this database to answer the following research questions: **Q1:** What are the temporal and biogeographical patterns of invasion from non-native human disease vector mosquitoes? **Q2:** What are the main introduction pathways and means of transportation for these mosquitoes, and their establishment success? **Q3:** Where did the mosquito species originate and where were they introduced to? **Q4:** What are the cold- and hotspots of mosquito introductions and establishment, and what is driving these patterns?

## Results

Of the 186 mosquito species acting as vectors for human pathogens^38^, we collected 698 first introduction records into new regions. These took place in 288 regions on all continents except for Antarctica. Of these, 610 records (i.e., 87.4 % of total) in 283 regions led to species establishment.

### High diversity of introduced non-native mosquitoes

45 mosquito species have been introduced to regions where they are non-native, accounting for 24.2% of all mosquito species known to transmit pathogens in the wild. Among these, 27 species have successfully established. These introduced species belong to five genera: *Aedes, Anopheles, Armigeres, Culex,* and *Mansonia*. Specifically, from the genus *Aedes* 15 species were introduced to a total of 468 regions, of which 10 species are established today in 408 regions. From the genus *Anopheles* likewise 15 species were introduced, but only in 35 regions, of which six species were able to establish in 16 regions. *Armigeres* has one introduced species with a single introduction and establishment record. Twelve species of the genus *Culex* have been introduced in 192 regions, of which nine species have become established in 184 regions. There are two introduced species of the genus *Mansonia* with two regional introduction records, of which one became established (Figure 2A, Suppl. Material Fig. 1, Suppl. Material Table 1).

**Figure 1.**
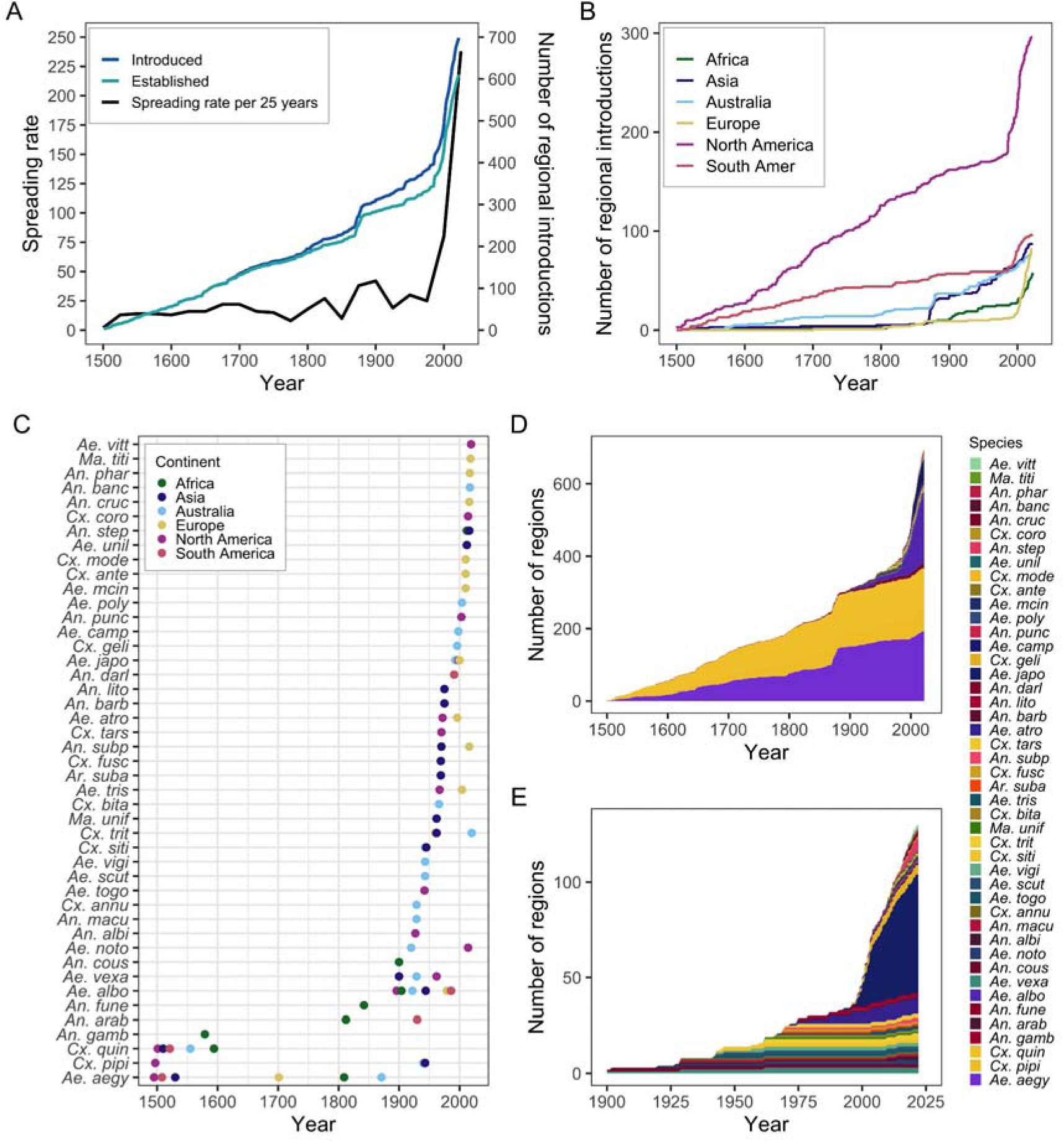
Temporal evolution in the cumulative number of first records of non-native mosquito species worldwide (A) and by continent (B). Spreading rates correspond to the total number of introduced species for a period of 25 years. Cumulative trends of first records at the global scale are shown for records regardless of species’ establishment in regions of introduction, as well as for records leading to species establishment. Timing of species’ first introduction records by continent (C). Cumulative timeline of the number of first introduction records for all species (D) and for species with global first record after 1900 (E). The timelines are shown for each of the 45 species (Names and abbreviations in Suppl. Material Table 6) with first introduction records in non-native regions. Figure legend shows species in descending order in terms of global first recording dates (more recently recorded species are showing first). Number of regional introduction records per continent can be found in Suppl. Material Table 3.

**Figure 2.**
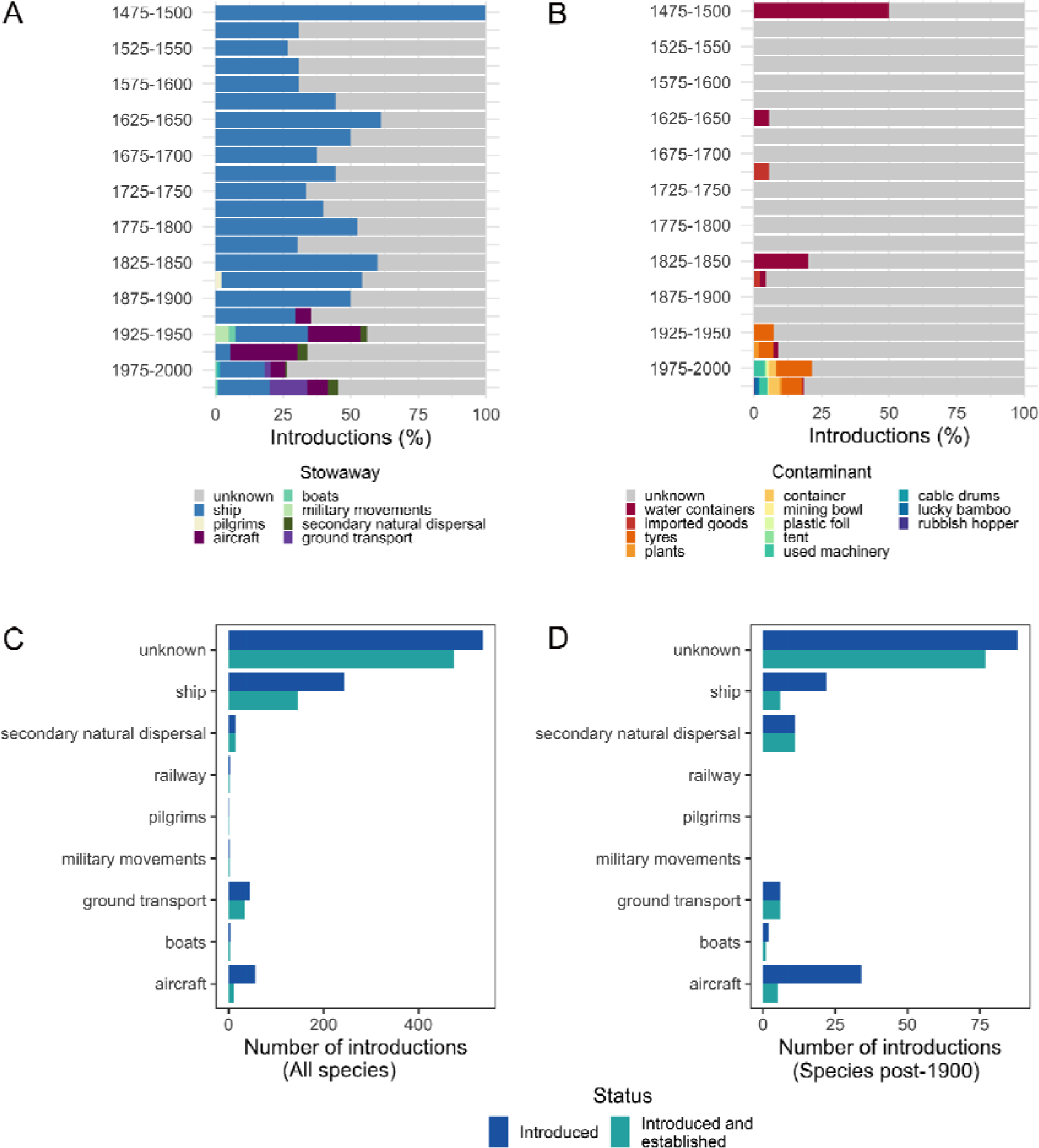
Proportion of transportation vectors (A) and transportation contaminants (B) in 25 year intervals of all introductions. Establishment success of the species depending on their transportation vector. (C) All species; (D) Species spreading post-1900.

**Table 1.**
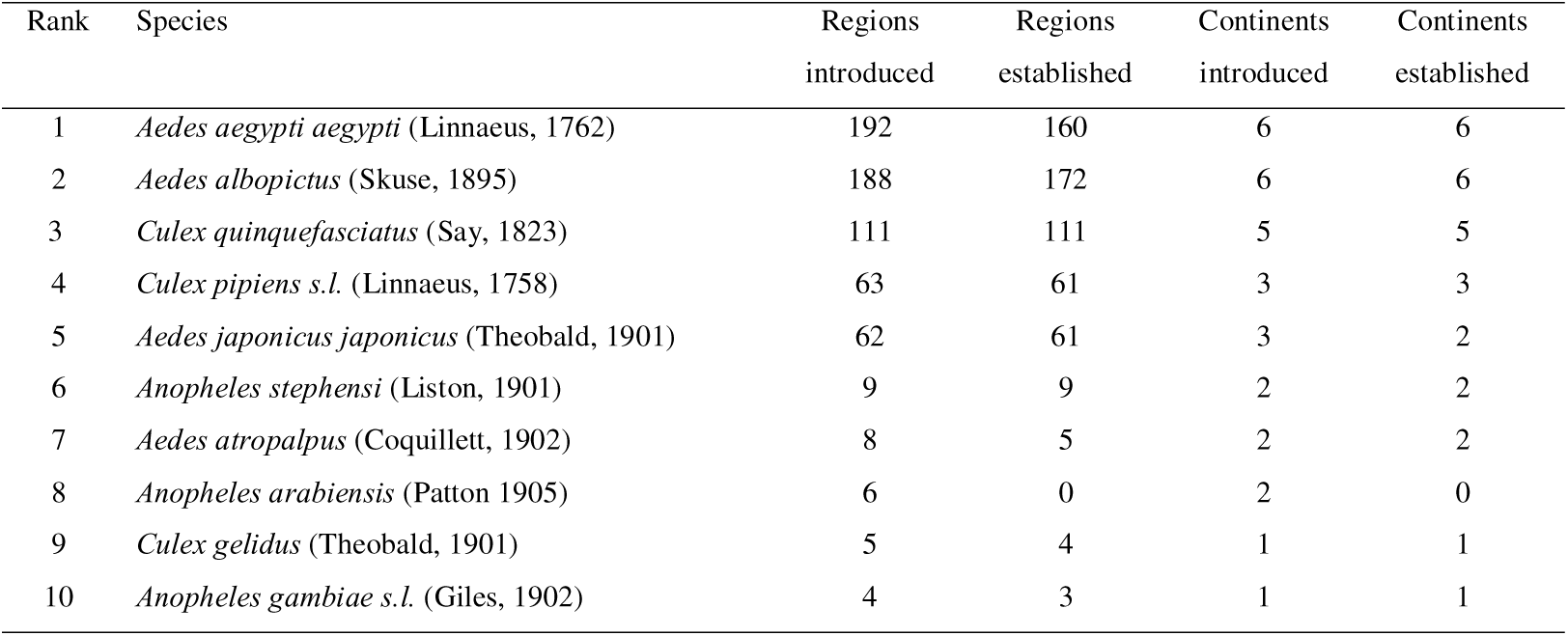
The ten most widely introduced species of the Global Non-Native Mosquito Database, with the total number of regions and continents where a given species was recorded.

| Rank | Species | Regions introduced | Regions established | Continents introduced | Continents established |
| --- | --- | --- | --- | --- | --- |
| 1 | *Aedes aegypti aegypti* (Linnaeus, 1762) | 192 | 160 | 6 | 6 |
| 2 | *Aedes albopictus* (Skuse, 1895) | 188 | 172 | 6 | 6 |
| 3 | *Culex quinquefasciatus* (Say, 1823) | 111 | 111 | 5 | 5 |
| 4 | *Culex pipiens s.l.* (Linnaeus, 1758) | 63 | 61 | 3 | 3 |
| 5 | *Aedes japonicus japonicus* (Theobald, 1901) | 62 | 61 | 3 | 2 |
| 6 | *Anopheles stephensi* (Liston, 1901) | 9 | 9 | 2 | 2 |
| 7 | *Aedes atropalpus* (Coquillett, 1902) | 8 | 5 | 2 | 2 |
| 8 | *Anopheles arabiensis* (Patton 1905) | 6 | 0 | 2 | 0 |
| 9 | *Culex gelidus* (Theobald, 1901) | 5 | 4 | 1 | 1 |
| 10 | *Anopheles gambiae s.l.* (Giles, 1902) | 4 | 3 | 1 | 1 |

The most widespread non-native vector mosquito species is *Ae. aegypti* recorded in 192 regions, followed by *Ae. albopictus* (188 regions) and *Cx. quinquefasciatus* (111 regions). *Aedes albopictus* also is the species with the highest number of regions (172) with established occurrences. Both, *Ae. aegypti* and *Ae. albopictus* have been introduced and established on all continents except for Antarctica. However, *Cx. quinquefasciatus* has not yet been introduced to Europe and Antarctica (Table 1).

### Pronounced increase in first records and species introduced since the mid-20^th^ century

The first known records of vector mosquitoes started with the beginning of intercontinental shipping in the 15^th^ century (Figure 1A). A substantial increase was then observed at the end of the 19^th^ century, with a rise in introductions mainly in Australia and Asia (Figure 1B). This was followed by a further increase in first detections from 1900 onwards, which may be partly due to more attention being paid to mosquitoes as their role as disease vectors was recognized in the same year^39^.

Since the mid-20^th^ century, there has been a rapid increase in newly recorded non-native mosquitoes, particularly in Europe, the Americas, and Africa. Nearly half (49 %) of all first records occurred after 1950 (n=342). Additionally, 28 of the 45 non-native vector mosquitoes were recorded for the first time after this date, and of these, 13 species were recorded for the first time after the year 2000 (Figure 1C-E). Concurrently, the volume of trade has increased 30-fold since 1950^40,41^, likely driving the rapid pace by causing an increased number of accidental introductions of, often small, inconspicuous taxa. This has been observed in ants^6^, plant pests^42^, earthworms^9^, and individual mosquito taxa^26^.

### Widening modes of transport and introduction

Most modes of transport and introduction of the species are unknown, but available data shows a shift in dominance from ship-based transportation to an increasing importance of aircrafts, ground-based transportation, and unaided secondary spread (Figure 2A). Known commodities carrying propagules of the species are highly varied (Figure 2B. The earliest report of mosquito introductions happened through standing water containers (water storage jars) on sailing vessels during slave trades at the end of the 15^th^ century^24^. Import via dried breeding sites in tyres appeared as an important contaminant in the 1900s. Plants were identified as notable pathways after 1950. Other commodities or objects such as containers, used machinery, a tent, rubbish bins, plastic foil, cable drums and mining bowls have been observed sporadically since the end of the 20th century^43^. Relevantly, while the role of aircraft in the introduction of species has become increasingly important, only a small proportion of these introductions establish in the new region (Figure 2C). Conversely, the relative importance of shipping as a pathway for the introduction of species has decreased since 1900 and the proportion of species that become established after transportation by ship has also decreased (Figure 2D).

### Many emerging non-native mosquitoes stay put, but a relevant fraction is spreading

We found that the number of regions where species have been recorded and the minimum residence time since their first global record are positively correlated (Pearson *r* = 0.604, p < 0.001 for records regardless of establishment and r = 0.609, p < 0.001 for those leading to establishment), indicating that species have been expanding their ranges over time. However, these correlations do not hold for species first recorded globally after 1900 and 1950 (Suppl. Material Fig. 5), suggesting that this finding is not true for emerging non-natives species. To further explore this relationship, we characterized the spread ranges of each species using a set of spatial variables and summarized them through projection into a Principal Component Analysis (PCA; Figure 3). Following this, k-means clustering was used to broadly classify species into four groups based on similarities of their spread ranges.

**Figure 3.**
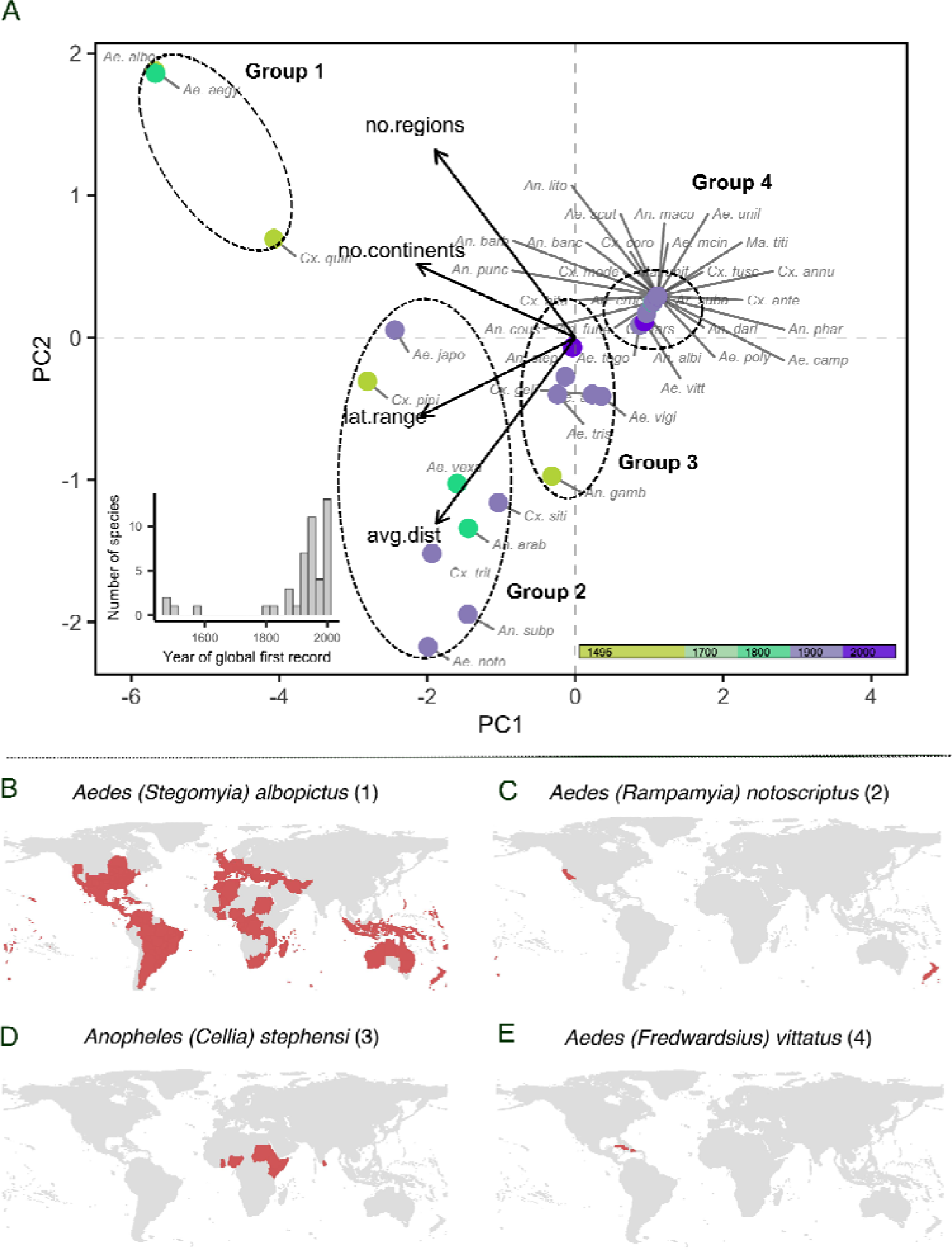
Principal Component Analysis (PCA) of species introduced outside native ranges over time. The axes of the PCA consider the number of regions the species was introduced (no.regions), number of continents (no.continents), latitudinal range between northernmost and southernmost region (lat.range) and average distance between all regions (avg.dist), where the x-axis explains 81.63 % and the y-axis explains 14.86 % of the total variance. Colors represent the timing of the global first record of each species. The inset histogram represents the number of global first records of vector mosquitoes per 25-year period. Maps below show the global distribution of first records (red) for an example species of each of the four groups identified.

The first group includes early cosmopolitan spreaders, comprising three well-known species: *Aedes aegypti, Ae. albopictus, and Cx. quinquefasciatus* (Group 1). While *Ae. aegypti* and *Cx. quinquefasciatus* began their global spread as early as the 15th century, *Ae. albopictus* only started spreading in 1896. These species are now established in numerous regions across most continents (Table 1). Another group consists of eight species: *Aedes japonicus, Ae. notoscriptus, Ae. vexans, An. arabiensis, An. subpictus, Cx. pipiens* s.l., *Cx. sitiens, Cx. tritaeniorhynchus* (Group 2), which either spread to many regions or overcame very large geographical distances in their spread. The third group contains species with medium wide non-native range sizes of up to nine regions colonized. Notably, five out of six species in this group began spreading in the 20^th^ century. These species include *Ae. atropalpus, Ae, triseriatus, Ae, vigilax, An. gambiae, An. stephensi* and *Cx. gelidus*. Finally, the most numerous group comprises 28 species having narrow non-native ranges. Of these, 22 were introduced only to a single non-native region. The remaining six species (namely: *Aedes togoi, Ae. vittatus, An. albimanus, An. coustani, An. funestus and Cx. tarsalis*) were introduced to two regions (Group 4). Most of these single-region introductions, representing 19 species (82.6% of the total), occurred after 1965.

### Dominant, but shifting roles of Africa and Asia in biogeographical flows to the rest of the world

#### Species flows across continents

The intercontinental flows of species show that Asia and Africa play a dominant role as donor continents of non-native mosquitoes (Figure 4A). Despite this, we also find a relevant number of species originating from North America (n=5), Australia (n=4), and the Old World (n=6) (i.e., Africa, Asia and Europe combined). In contrast, Europe and South America primarily are recipients of introduced mosquitoes. In Australia, there is predominantly an intracontinental flow of introduced species, and introductions from Asia. Among continents, Australia, Europe, and North America had the highest number of regional introductions (Table 2, Figure 4).

**Figure 4.**
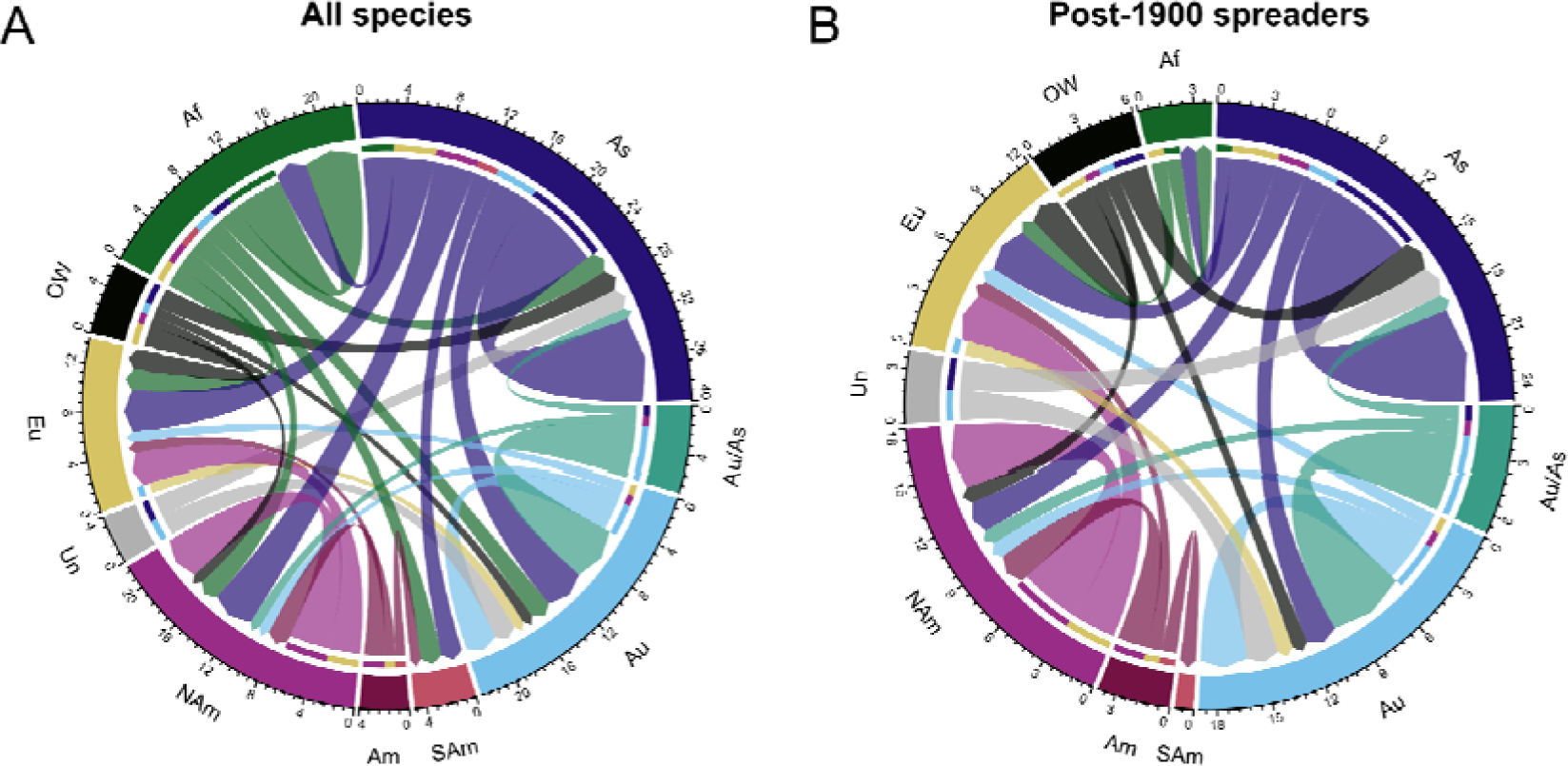
Global species flows of non-native mosquitoes, representing the introduced species richness to and from each continent. Each species recorded on a continent was counted only once. (A) Flows for all species. (B) flows for species that started spreading after 1900. Af: Africa, Am: America, As: Asia, Au/As: Australia and Asia, Au: Australia, Eu: Europe, NAm: North America, OW: Old World (i.e., Africa, Asian and Europe combined), SAm: South America, Un: Unknown

**Table 2.**
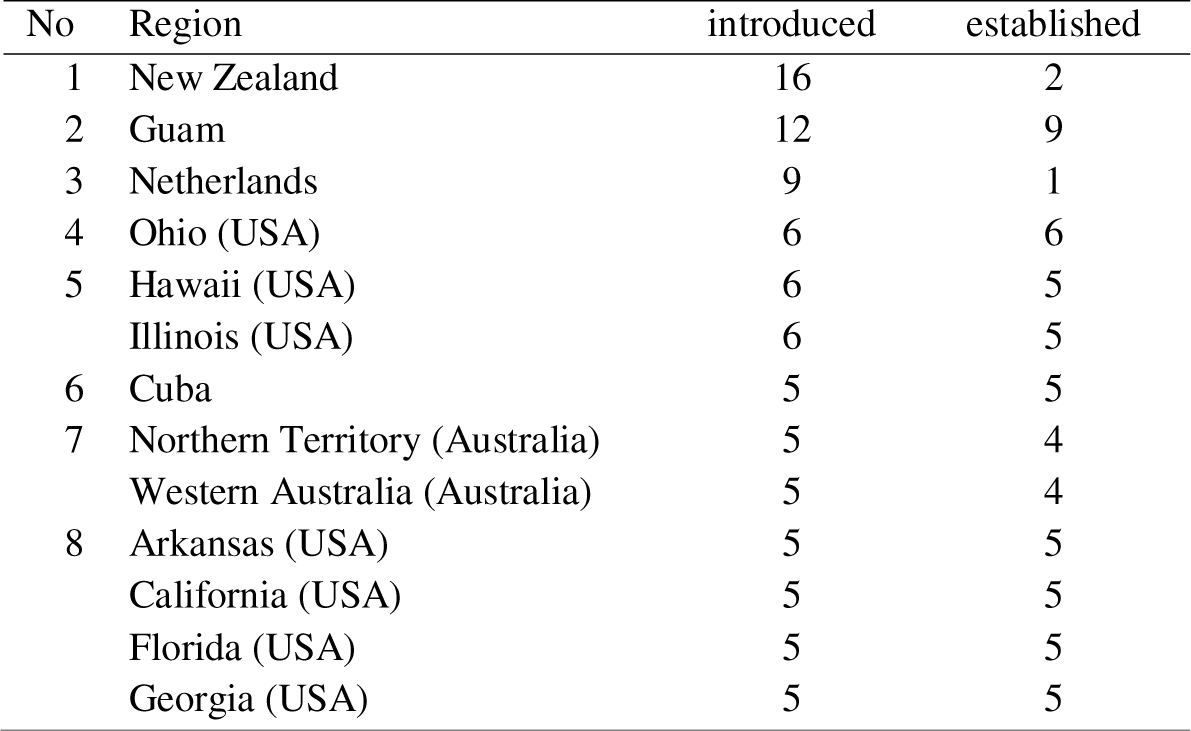
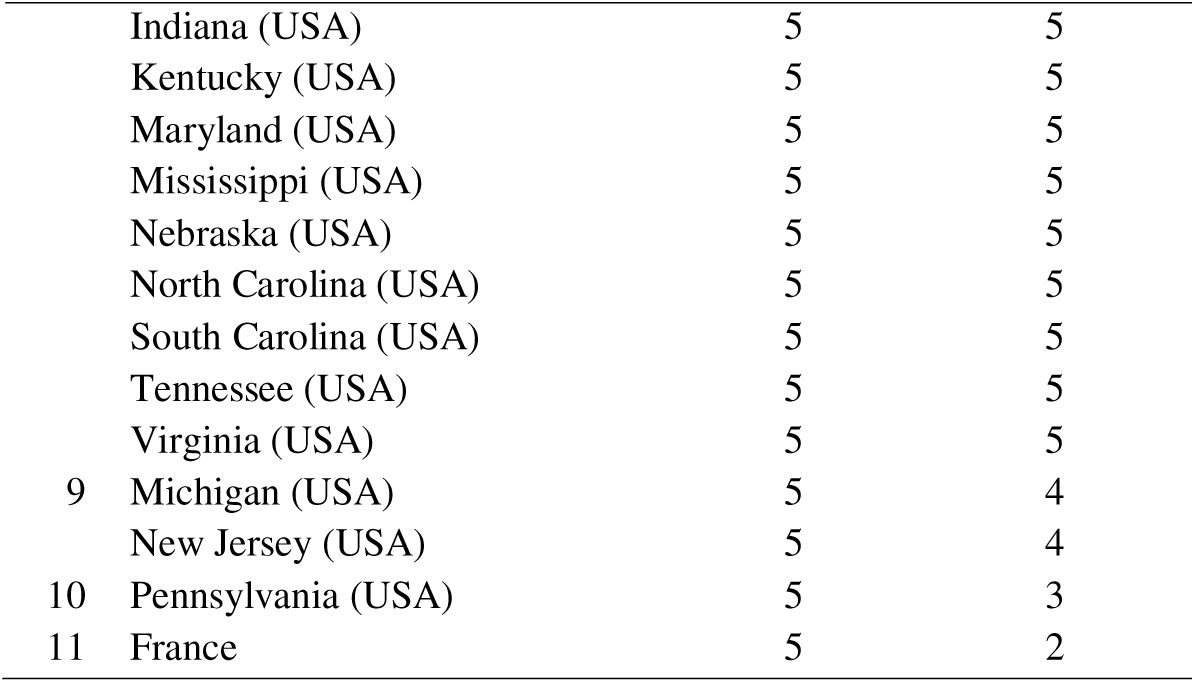
Regions with five or more introductions of non-native vector mosquito species.

#### Post-1900 colonization trends

When focusing on species that started to spread from 1900 onwards, we found that Africa’s role as a donor continent is reduced and that most introduced species come from Asia, Australia, and the Americas. Africa receives only one additional mosquito species from outside the continent, and only two species are beginning to spread from there. All species native from Australia and the Americas only began to spread after 1900, either within their native continent or to Europe. A single European species (*An. maculipennis*) was introduced to only a single region, New Zealand (Figure 4B).

### Introduction numbers are higher in Australia, Europe and North America

Of the 477 assessed regions, 288 have records of the introduction of at least one non-native vector mosquito, representing 63.4% of the world’s land area. New Zealand had the highest number of recorded first introductions (n=16), followed by Guam (n=12), the Netherlands (n=9), and three U.S. states (Hawaii, Illinois, Ohio) (n=6) (Table 2, Suppl. Material Fig. 6). The majority of regions, 62.2%, recorded the introduction of only one or two new species. Guam tops the ranking for established non-native species (n=9), followed by Ohio (n=6), and 15 eastern U.S. states, California, and Cuba (n=5). Despite a strong correlation between the introduction and establishment of species (Pearson r = 0.76, p < 0.001), some regions exhibit large differences between the species richness of introductions and establishments, such as New Zealand, the Netherlands, France, Guam, Mauritius, Reunion, Pennsylvania, Barbados, Belgium, and the UK.

We identified hot- and coldspots of introduction, defined as countries where the recorded number of introductions is higher or lower than expected, respectively. The hotspots include New Zealand, the Netherlands, USA, Mauritius, France, Palau and Italy (Figure 5A), while the Cook Islands, Norway, Poland, Ukraine and Finland were identified as coldspots. Focusing on species that started to spread only after 1900, we identified hotspots of introduction in New Zealand, the Netherlands, USA, France, Cuba, and Italy (Figure 5B). The United Kingdom, Norway, Poland, Portugal, Austria, and Ukraine were identified as coldspots for species spreading after 1900. Compared to hotspots of establishment, the introduction hotspots New Zealand, France, and the Netherlands performed better than expected (Figure 5C,D). On the contrary Australia is one of the hotspots of species establishment for all species, but this has changed for species that spread after 1900.

**Figure 5.**
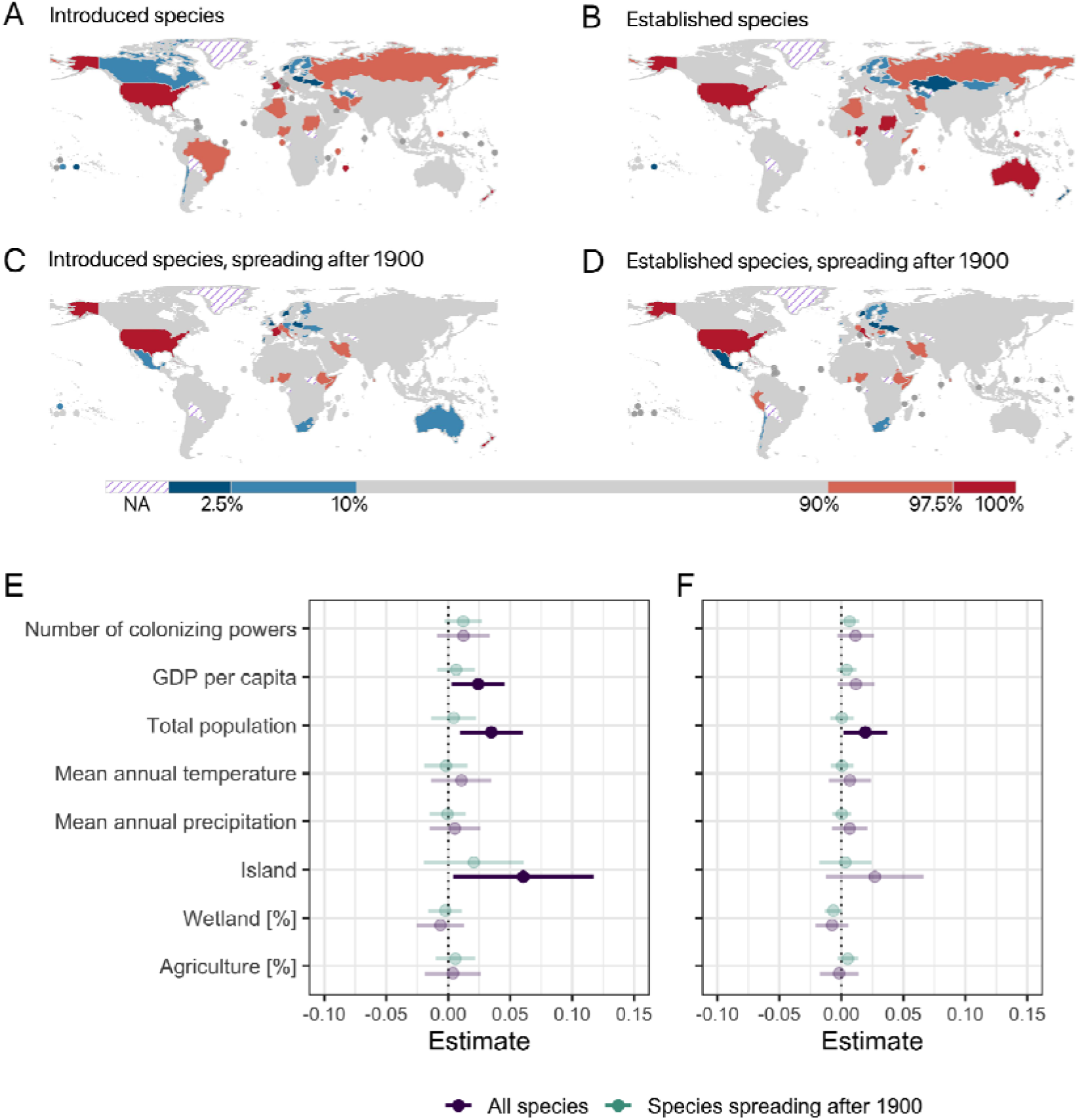
Country level cold and hotspots of introduction (left column) and establishment (right column). Maps represent cold- and hotspots of all species (A,B) and for species spreading post-1900 (C,D). Colors indicate countries with higher (red) and lower (blue) vector mosquito numbers than expected. Note the non-linear scale of the color legend. For countries in purple stripes IAS measures are unknown. Countries smaller than 27,106 km² (equivalent to Haiti) are shown as circles. Results of regression models testing the predictors of socio-economic and environmental variables for hot- and coldspots of mosquito invasion. Effect sizes for eight variables tested in GLMMs for cold- and hotspots of species introduction (E) and establishment (F). Purple color shows model results for all species, turquoise for models with species introduced in non-native regions only after 1900. Darker colors indicate significant effects in the models (p < 0.05).

### GDPpc and Population size show highest importance for mosquito species introduction

The multivariate linear mixed model analysis to identify drivers for the variation in cold- and hotspots of mosquito introduction and establishment, revealed that GDP per capita is significantly positively associated (p < 0.05) with the propensity of countries to receive species introductions (Figure 5E). The total population size of a country had a highly significant effect (p < 0.01) on species introductions, and a significant effect (p < 0.05) on species establishment (Figure 5F). Insularity also had a significant effect (p < 0.05) on the extent to which species were introduced.

## Discussion

We compiled and analyzed a comprehensive global database of first records of mosquito vectors for human diseases across regions and show that a large and increasing number of mosquito species are being dispersed by human activity, following trends in the globalization of trade and transportation. Additionally, we observed substantial shifts in the region of origin of species from Africa to Asia over time and the pathways through which they are introduced, but fairly stable patterns in their recipient regions (i.e., the Americas, Australia and Europe).

We show that the rates of mosquito introductions into new regions and the emergence of novel non-native species have been increasing in recent centuries, with a particularly pronounced increase observed in recent decades. While it is plausible that enhanced surveillance and import controls have contributed to this trend by increasing detection rates (e.g., at ports and airports^43^), the dramatic rise in first records of newly introduced species strongly matches the substantial expansion and diversification of global trade during the mid-19^th^ century^41^. During this period the frequency and volume of commodities transported across continents increased and travel times decreased substantially^41^. Furthermore, the advent of rapid transportation methods, such as cargo planes, has enabled the unintentional dissemination of short-lived propagules, including adult mosquitoes^29,44^. Collectively, these factors are increasing the opportunities for the unintentional spread of mosquito vectors and are likely to continue doing so in the coming decades ^25,45,46^.

Most introduction records lead to establishment. This very high proportion of established introductions suggests that there may be many casual introductions, which remain unnoticed. On the other hand, many species emerging after 1950 have not become established (yet), with 12 out of 28 species (43%) failing to establish in a new region. This low establishment rate may be due to enhanced surveillance and import controls that may detect and intercept these species at the introduction stages or in circumstances that would not typically lead to establishment, such as single adult male mosquitoes^24^. Additionally, the low establishment rate may reflect lag effects in the reporting of the species or between the introduction of a species and its rapid population growth and spread where detection becomes more likely^47^. This lag phase can vary greatly among species and environments, lasting from a few years to several decades^48^. Therefore, while some detected species are not currently established, they could become so in the future.

A species’ establishment success is also influenced by its habitat requirements and adaptability, including the availability of suitable oviposition sites^49^. For example, the genus *Aedes* exhibits approximately 2.5 times more introduction records than the genus *Culex* and over 13 times more than the genus *Anopheles*. This high number of introductions for *Aedes* can be attributed to the ability of their eggs to withstand desiccation, allowing them to be transported over long distances and extended periods (3-6 months) in both artificial and natural carriers that do not require standing water^50^. In contrast, the eggs of other genera, lacking this desiccation resistance, often perish when transported in containers that dry up during transit, leading to unsuccessful establishment in new regions and inherently reducing their potential for further dispersal.

Historical changes in global trade and travel dynamics have led to pronounced shifts in the donor regions of species being transported, most notably a shift from African-origin species to those from Asia. Despite these shifts in geographical provenance, we found that the hotspots of introduction have remained rather stable. These include primarily the USA, several Central European and tropical African countries, and New Zealand, with the European, Australian, and North American continents also consistently having high ratios as destinations.

We also found that socioeconomic variables, in particular per capita GDP and population size, have a significant positive relationship with the invasion and establishment success of mosquitoes, as has been found for other taxa^17^. In contrast, colonial relationships, which are a major factor for the introduction of other non-native species such as plants (including intentionally introduced crops)^5^, were not found to have a significant effect on the introduction of mosquito species, possibly because they were always introduced unintentionally. Insularity also had a significant effect on the extent to which species were introduced, consistent with findings for other taxa^17^. These results highlight the necessity of incorporating socio-economic relationships into invasion prevention strategies to address the complexities of global flows of commodities and transportation vectors susceptible to carrying mosquito propagules. Research on mosquito spread should expand therefore beyond the widely explored effects of climatic variables and their changes^51,52^, and include land-use changes and socio-economic variables such as poverty, trade and travel^53^.

Ultimately, our work shows that increased attention should be paid not only to well-known primary vectors such as *Ae. albopictus* and *Ae. aegypti*, but also to other increasingly dispersed mosquito vectors, as they can significantly contribute to the maintenance of pathogens^54^. The adaptability of mosquito species^55^ and pathogens^56^, combined with the escalating rate of introductions, poses a considerable challenge for global public health. Mitigation and prevention efforts should focus on pathways of introduction, prioritize source and target regions identified as hotspots of non-native mosquitoes, and implement rapid response protocols following early detection. These strategies are essential to mitigating the impact of non-native mosquito species on public health in an increasingly interconnected world.

## Methods

### Data compilation

We systematically searched the literature for records of human-introduced non-native mosquito species. For the standardization of species names, we used the Walter Reed Biosystematics Unit (WRBU) taxonomic reference which also lists synonyms for each species^57^. Only mosquito species known to be vectors for human diseases in the wild (n=186) were considered (i.e., naturally occurring with a human pathogen). This condition must be confirmed either by at least two scientific articles or one source together with an entry in the mosquito catalogue^57^ that confirmed the link to a disease. We searched for scientific publications on Google Scholar between March and December 2023, using all combinations of the following search terms: “*species name*” AND “*country name*” AND “introduction” OR “invasion” OR “first record”. Additionally, we reviewed existing literature reviews^e.g., 24,36,57^ and citing documents. We compiled the results in a database, in which each row represents a single record of the first record of a non-native mosquito species in a new region. In case the species was subsequently eradicated or went extinct, but was again reintroduced later, another new record was created. We only considered records resulting from human mediated introductions or the natural secondary spread of populations originating from them (i.e., so-called non-native species^2^). To standardize the record collection, we used regions consisting of countries and added the first-order administrative divisions of the six largest countries (Australia, Brazil, Canada, China, Russia, USA) as obtained from GADM (“https://gadm.org/”). Additionally, certain islands such as Corsica, Sicily, Sardinia, Crete, Madeira, the Azores, the Galapagos Islands, the Balearic Islands, the Canary Islands and the Torres Strait are considered separately, as water is a barrier to the migration of mosquito species; in total, this approach resulted in a layer of 477 regions.

In a few cases, we could not be certain of the reliability of introduction records. For example, in some situations, it was unclear whether the species’ appearance in a region was due to natural dispersal from native populations or human-mediated introduction. Therefore, records for which reliability was uncertain were not included in our database.

The database contains entries on the names of the species, the year of first record, the mode of transportation and introduction (when available), the status of establishment and the source of the information. For some introductions that date back before the 20th century, we used approximations of the date of introduction based on disease descriptions and estimations from other sources (e.g. *Ae. aegypti* reports of yellow fever etc.).

### Introduction pathways

The mode of introduction of species is distinguished between two main classes: dispersal vectors (transportation stowaways) and the commodities serving as carriers during dispersal (transportation contaminants). We categorized these two large classes into more specific groups. For transportation stowaways we considered ships, boats, ground transportation (cars), trains, aircraft, secondary natural dispersal (after human introduction to the continent), and unknown sources. For transportation contaminants we considered water containers, plants, tires, lucky bamboo, used machinery, containers, and imported goods and objects. Introduction records were segmented into 25-year intervals, and the proportions of stowaways and contaminants for each period were presented.

### Assessing similarities in non-native species ranges

Principal Component Analysis (PCA) was employed to visually explore (dis)similarities in the non-native ranges of species. The variables describing each species included: 1. The number of regions where the species was recorded. 2. The number of continents where the species was recorded. 3. The average distance (in km) between centroids of the regions where the species was recorded. 4. The latitudinal range, defined as the distance (in km) between the highest and lowest latitude of the centroid of the region where the species was recorded. Following species projection into a bidimensional PCA, whose axes explained 96.48% of total variance in the data, we used the k-means algorithm to group species according to similarities in their spread ranges, and the elbow method to identify the optimal number of clusters (Suppl. Material Fig. 3 and 4)^58^. To perform the PCA, we used the ‘prcomp()’ function and the k-means was implemented using the ‘kmeans()’ function both from the *base* R package. For visualization and cluster analysis of the PCA results we used the ‘fviz_pca_biplot()’ function from the *factoextra* package (version 1.0.7)^59^.

### Continental flows

To visualize the exchange between continental regions, we linked the native region(s) of each species to the continent(s) of its non-native area of introduction. First, we identified the native continent of each individual species according to the current state of knowledge in the literature (Suppl. Material Tab. 4). For a few species, the native ranges were not limited to a single continent. Species mentioned to be native to more than one continent among Europe, Africa, and Asia were grouped under the term ‘Old World’. Species whose native range was even less precisely defined (e.g. Old World plus Australia) were grouped in the ‘unknown’ category. In some cases, we grouped two continents (i.e., Americas, Australia and Asia). The delineation of continents was made according to the Suppl. Materials Fig. 2. We calculated the number of species introduced to each continent, with each species introduced to a continent counted only once and represented by the arrows pointing to the continents. This included both intercontinental and intracontinental introductions, accounting for instances where a species was introduced to a new region within the same continent. Flow diagrams were created using the *circlize* package (version 0.4.15)^60^.

### Assessing patterns and drivers of species distributions

To assess the geographical patterns of species introductions and establishment, we summed the number (richness) of first records per region and the number of species that successfully established. To assess possible temporal shifts in the patterns of introduction and establishment, this was performed considering 1) all species and 2) only those that had their first global record after 1900. To identify hotspots and coldspots of species introduction, we used a General Linear Mixed Model (GLMM^61^ in which we used the number of first records as dependent variable and as explanatory variables we used country area size and a proxy of country’s recording effort. The first variable was used to account for species-area relationships, wherein regions with larger areas tend to harbor more species^62^. The proxy variable for recording effort encompassed both proactive measures (e.g., surveillance initiatives) and reactive responses (e.g., national action plans for invasive species management) as defined by Early et al.^63^. These authors ranked countries based on their capacity to prevent the introduction of Invasive Alien Species (IAS) and their ability to identify and control emerging invasions. This ranking ranged from 0 (lowest capacity) to 3 (highest capacity) and was derived from reports to the Convention on Biological Diversity. Factors considered in the ranking included existing legislation on cargo inspection procedures, public awareness programs, and resources allocated for species identification and reporting. We used the average between proactive and reactive effort. Six countries are at the top of the ranking: Australia, Austria, Canada, New Zealand, the United Kingdom, and the USA. This variable is available at the country level. Consequently, the models were developed at this spatial resolution after aggregating first record data from sub-national divisions. Due to limitations in the spatial coverage of the recording effort proxy, 47 countries had to be excluded from the analysis. The final models encompassed a total of 181 countries. We fitted the models using the *glmmTMB* package^61^, assuming a negative binomial distribution of errors and using continents as a random effect to account for spatial autocorrelation among regions. The residuals of these models were then ranked to calculate the upper and lower quantiles, which serve as boundaries for defining hot- and coldspots for the introduction and establishment of non-native mosquitoes (i.e. upper/lower 10% and 2.5%).

To explore the factors driving the geography of hot- and coldspots, we used a GLMM to relate the residual values (log-transformed) to a set of country-level socioeconomic and environmental variables. The variables were (1) gross domestic product per capita (GDP), (2) total trade openness in US$, (3) time of occupation by one of the eight main European colonial empires (4) Number of European empires that had a colonial influence on the country, (5) total human population, (6) long-term average yearly temperature, (7) long-term average yearly sum of precipitation, (8) percentage of land cover by wetlands and (9) agriculture, (10) insularity (‘yes’, ‘no’), and (11) the absolute latitude. A detailed list of these variables, along with their measurement units and respective data sources, is available in Suppl. Material Tab. 4.

Of these variables, trade openness and occupation by colonial empires have a more intricate meaning and require a deeper explanation. Trade openness is a measure of a country’s trade globalization. This measure was calculated as an average between 2017 and 2021 of imports and exports of total trade in goods and services as a percentage of the country’s GDP^64^. To represent colonization patterns, we utilized the Empire Database^5^, which originally had data for the four main colonial empires, but was meanwhile extended to eight empires (unpublished data). This dataset provides information on the duration of occupation by the eight European colonial empires: British, Dutch, Spanish, Portuguese, and extended with Belgian, French, German and Italian. We calculated the total duration of occupation for each country and the number of colonial powers that occupied each country.

To avoid redundancy between explanatory variables, we only included those with pairwise Pearson correlation coefficients below |0.7| and a variance inflation factor below 3, calculated using the *usdm* package (version 2.1.7)^65^. The variable “total trade openness in US$” was highly correlated with GDP per capita and population size. Similarly, “absolute latitude” was correlated with temperature, and “colonial occupation time” was correlated with the number of colonial empires. Consequently, these three variables were excluded from the models. Several variables (see Suppl. Material Tab. 4) were log transformed to ensure normality and numeric variables were scaled to make them comparable^66^. The correlation matrix for the predictors used in the models to test the drivers can be found in the Suppl. Material Fig. 7. The model assumed a gaussian distribution of errors.

All calculations were done using R version 4.3.2 (2023-10-31)^67^ in RStudio^68^. All figures were created using the *ggplot2* package (version 3.4.4)^69^. All maps were produced using QGIS (version 3.30.1-’s-Hertogenbosch)^70^.

## Acknowledgements

We gratefully acknowledge the support of the Portuguese Foundation for Science and Technology (FCT) for funds to GHTM - UIDB/04413/2020 & UIDP/04413/2020 and LA-REAL – LA/P/0117/2020. RP acknowledges support from FCT (grand no. PRT/BD/153694/2021). CC and JLZ acknowledge support from FCT through support to CEG/IGOT Research Unit (UIDB/00295/2020 and UIDP/00295/2020). FE, BL and AG-R acknowledge funding from the Austrian Science Foundation FWF (grant no. I-5825-B). AS and DL acknowledge funding from the Austrian Science Foundation FWF (grant no. P 34688-B).

## Notes

### Competing Interest Statement

The authors have declared no competing interest.

