## Supplementary_Material_1 for "Global invasion patterns and dynamics of disease vector mosquitoes"

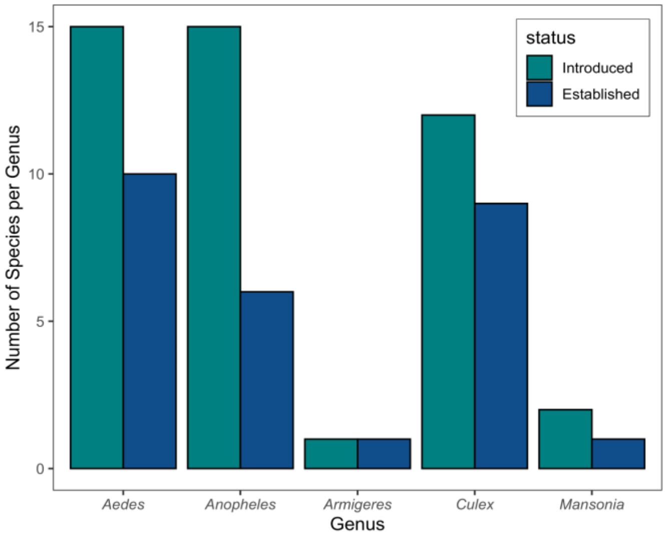


**Suppl. Material Fig. 1:** Number of human disease-transmitting mosquito species introduced by humans outside their native range compared to the number of species that have become established. Values are shown for genera.

**Suppl. Material Tab. 1:** Number of regional introduction records and number of species of each genus for introduced species as well as introduced and established species.

| **Genus** | **Regional introduction records** | | **Number of species** | |
|---|---|---|---|---|
| | **Introduced** | **Established** | **Introduced** | **Established** |
| ***Aedes*** | 468 | 408 | 15 | 10 |
| ***Anopheles*** | 35 | 16 | 15 | 6 |
| ***Armigeres*** | 1 | 1 | 1 | 1 |
| ***Culex*** | 192 | 184 | 12 | 9 |
| ***Mansonia*** | 2 | 1 | 2 | 1 |

**Suppl. Material Table 2:** Numbers of non-native mosquito species for donor and receiving continents.

| Continent | All species | | | Species started spreading after 1900 | | |
|---|---|---|---|---|---|---|
| | Number of non-native species native from this continent (total: 45) | Number of species introduced to this continent | Number of species introduced to this continent from outside the continent* | Number of species originated from this continent (total: 38) | Number of species introduced to this continent | Number of species introduced to this continent from outside the continent* |
| **Africa** | 7 | 8 | 3 | 2 | 2 | 1 |
| **Asia** | 12 | 15 | 2 | 10 | 11 | - |
| Asia/Australia | 5 | - | - | 5 | - | - |
| **Australia** | 4 | 18 | 9 | 4 | 14 | 4 |
| **Europe** | 1 | 13 | 11 | 1 | 11 | 9 |
| Americas | 4 | - | - | 4 | - | - |
| **North America** | 5 | 15 | 9 | 5 | 11 | 5 |
| **South America** | - | 5 | 4 | - | 1 | - |
| Old World | 4 | - | - | 4 | - | - |
| uncertain | 3 | - | - | 3 | - | - |

*Species of uncertain origin, along with species introduced to Asia from the Old World, and species originating from Asia and Australia introduced to one of these continents are not included in the count.

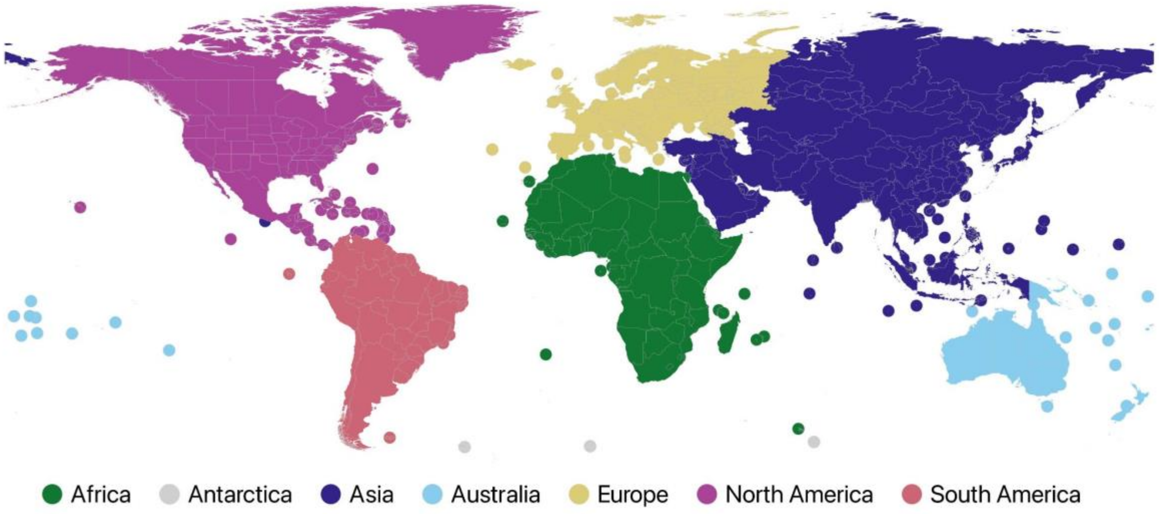


**Suppl. Material Fig. 2.** Delineation of continents (colors) used for the analysis in this paper. Delineation follows the seven continent model[1]

**Suppl. Material Table 3.** Number of regions non-native mosquito species were recorded after they were introduced to a continent. Colors represent the year of the first continental record. () represent the total number of regions in that continent. o 1495-1700 o 1700-1800 o 1800-1900 o 1900-2000 o 2000-2022

| Species | Number of introduced regions per continent | | | | | | SUM |
|---|---|---|---|---|---|---|---|
| | Africa (60) | Asia (126) | Australia (32) | Europe (115) | North America (105) | South America (42) | |
| Ae. aegy | 8 | 41 | 17 | 15 | 71 | 40 | 191 |
| Ae. albo | 27 | 17 | 13 | 36 | 59 | 36 | 188 |
| Ae. atro | | | | 4 | 4 | | 8 |
| Ae. camp | | | 1 | | | | 1 |
| Ae. japo | | | 1 | 18 | 43 | | 62 |
| Ae. mcin | | | | 1 | | | 1 |
| Ae. noto | | | 1 | | 1 | | 2 |
| Ae. poly | | | 1 | | | | 1 |
| Ae. scut | | | 1 | | | | 1 |
| Ae. togo | | | | | 2 | | 2 |
| Ae. tris | | | | 1 | 1 | | 2 |
| Ae. unil | | 1 | | | | | 1 |
| Ae. vexa | | 1 | 1 | | 1 | | 3 |
| Ae. vigi | | | 2 | | | | 2 |
| Ae. vitt | | | | | 2 | | 2 |
| An. albi | | | | | 2 | | 2 |
| An. arab | 4 | | | | | 2 | 6 |
| An. banc | | | 1 | | | | 1 |
| An. barb | | 1 | | | | | 1 |
| An. cous | 2 | | | | | | 2 |
| An. cruc | | | | 1 | | | 1 |
| An. darl | | | | | | 1 | 1 |
| An. fune | 2 | | | | | | 2 |
| An. gamb | 4 | | | | | | 4 |
| An. lito | | 1 | | | | | 1 |
| An. macu | | | 1 | | | | 1 |
| An. phar | | | | 1 | | | 1 |
| An. punc | | | | | 1 | | 1 |
| An. step | 8 | 1 | | | | | 9 |
| An. subp | | 1 | | 1 | | | 2 |
| Ar. suba | | 1 | | | | | 1 |
| Cx. annu | | | 1 | | | | 1 |
| Cx. ante | | | | 1 | | | 1 |
| Cx. bita | | | 1 | | | | 1 |
| Cx. coro | | | | | 1 | | 1 |
| Cx. fusc | | 1 | | | | | 1 |
| Cx. geli | | | 5 | | | | 1 |
| Cx. mode | | | | 1 | | | 1 |
| Cx. pipi | | 11 | 6 | | 46 | | 63 |
| Cx. quin | 3 | 8 | 22 | | 60 | 18 | 111 |
| Cx. siti | | 1 | 1 | | | | 2 |
| Cx. tars | | | | | 2 | | 2 |
| Cx. trit | | 1 | 1 | 1 | | | 3 |
| Ma. titi | | | | 1 | | | 1 |
| Ma. unif | | 1 | | | | | 1 |

**Suppl. Material Table 4.** Description and sources of the socio-economic, geographic and environmental variables that were considered as predictors used in the models to estimate the residuals and the models to explain the residuals.

| Variable | Description | Source |
|---|---|---|
| **Socioeconomic variables** | | |
| Gross domestic product (GDP) per capita (log transformed) | 5 year average between 2017 and 2021 | Worldbank (databank.worldbank.org)[2] |
| Total trade openness (log transformed) | 5 year average between 2017 and 2021, average of import and export in US$ at current prices in millions | UNCTAD dataset (United Nation Conference on Trade and Development)[3] |
| Colonial impact | Occupation time by the eight European colonial powers: British, Dutch, Spanish, Portuguese, Belgian, French, German, and Italian.<br><br>Number of European colonial powers that occupied the country. | Lenzner et al. 2023[4] and Lenzner (unpublished) |
| Human population density in the year 2021 (log transformed) | Number of individuals per km2 in the year 2021 | Worldbank (databank.worldbank.org)[2] |
| Sampling bias | the average between a country's proactive and reactive capacity towards invasive alien species | Early et al. 2016[5] |
| **Geographic variables** | | |
| Insularity | 1 = island; 0 = mainland | No specific source |
| Latitude of centroid | absolute value of decimal degree | Calculated in QGIS |
| Area (log transformed) | size of the region/country in km2 | Calculated in QGIS |
| **Environmental variables** | | |
| Average annual temperature | The average annual temperature from 1970 to 2000 in °C, resolution 30 Arcsec | worldclim[6] |
| Average annual precipitation | The average annual total precipitation from 1970 to 2000 in millimeters, resolution 30 Arcsec | worldclim[6] |
| Landcover (log transformed) | Copernicus Global Land Service: Land Cover 100m; percentage of wetlands/rivers and agricultural land | Copernicus[7] |

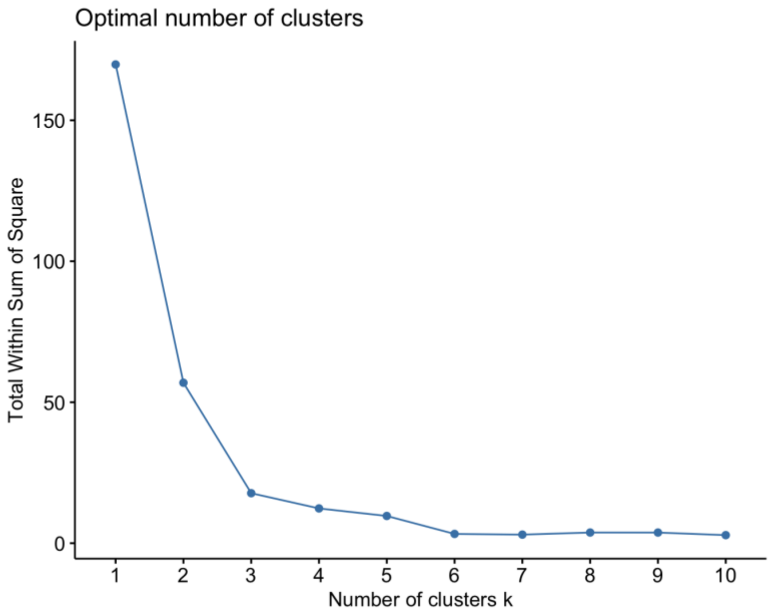


**Suppl. Material Fig. 3.** Elbow method to identify optimum number of clusters.

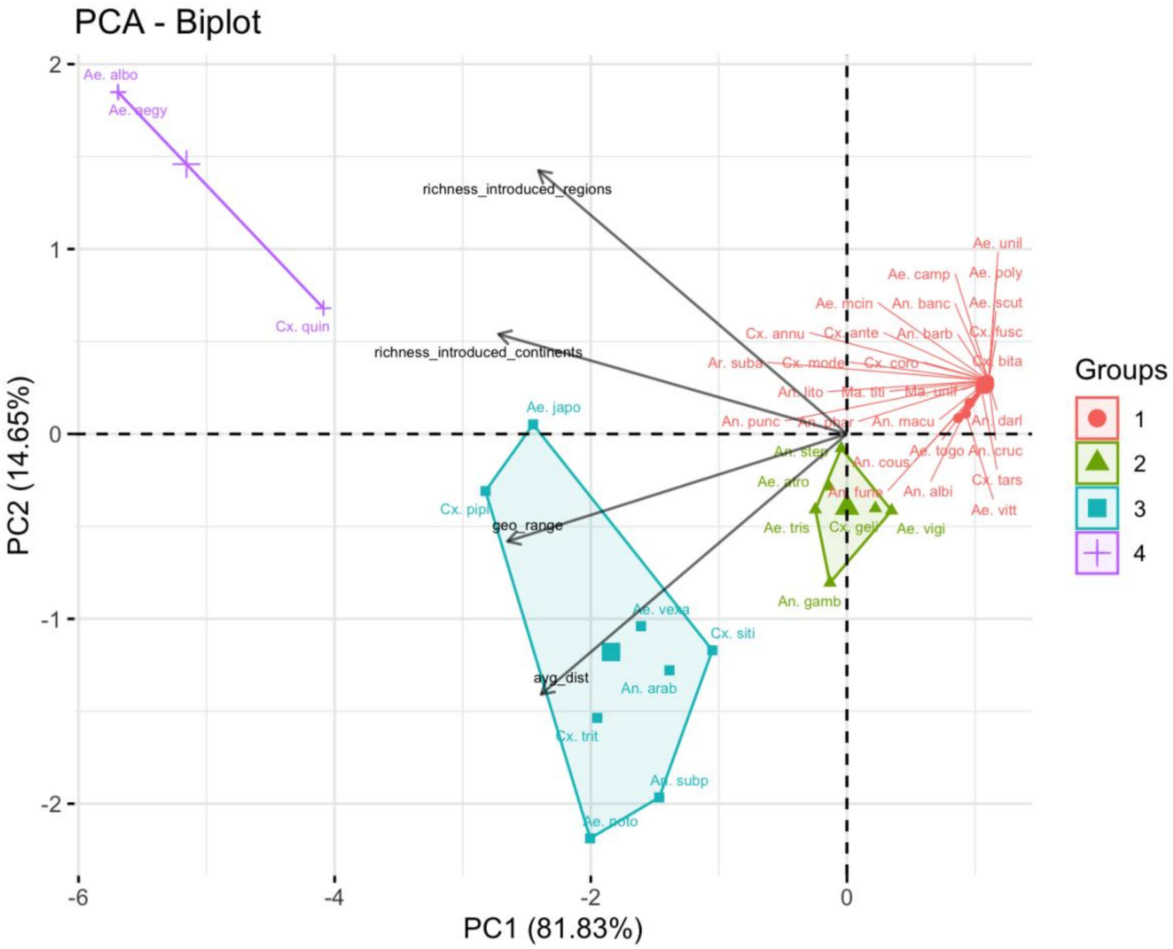


**Suppl. Material Fig. 4.** Biplot with 4 groups.

**Suppl. Material Tab. 5.** Assignment of native ranges of non-native species to continents and relevant data sources.

| *Species* | Native Continent | Native continentsif unclear | Source | Source Link |
|---|---|---|---|---|
| *Aedes (Aedimorphus) vexans* | Asia/ Australia | NA | Wilkerson et al. 2021 | https://doi.org/10.1353/book.79680 |
| *Aedes (Fredwardsius) vittatus* | Old World | Africa/Asia - Trop/Europe | Wilkerson et al. 2021; Díez-Fernández et al. 2018 | https://doi.org/10.1353/book.79680; https://doi.org/10.1186/s13071-018-2879-4 |
| *Aedes (Georgecraigius) atropalpus* | North America | NA | Medlock et al. 2015 | https://doi.org/10.1017/S0007485315000103 |
| *Aedes (Hulecoeteomyia) japonicus japonicus* | Asia | NA | Medlock et al. 2015; Tanaka et al. 1979 | https://doi.org/10.1017/S0007485315000103; https://mosquito-taxonomic-inventory.myspecies.info/sites/mosquito-taxonomic-inventory.info/files/Tanaka%20et%20al%201979.pdf |
| *Aedes (Ochlerotatus) camptorhynchus* | Australia | NA | Hearnden 1999 | https://www.researchgate.net/profile/Mark-Hearnden/publication/254094793_A_Health_Risk_Assessment_for_the_Establishment_of_The_Exotic_Mosquitoes_Aedes_camptorhynchus_and_Culex_australicus_in_Napier_New_Zealand/links/5473b5bb0cf245eb436db886/A-Health-Risk-Assessment-for-the-Establishment-of-The-Exotic-Mosquitoes-Aedes-camptorhynchus-and-Culex-australicus-in-Napier-New-Zealand.pdf |
| *Aedes (Ochlerotatus) mcintoshi* | Australia | NA | Wilkerson et al. 2021 | https://doi.org/10.1353/book.79680 |
| *Aedes (Ochlerotatus) vigilax* | Asia/ Australia | NA | Wilkerson et al. 2021 | https://doi.org/10.1353/book.79680 |
| *Aedes (Protomacleaya) triseriatus* | North America | NA | Medlock et al. 2012 | https://doi.org/10.1089/vbz.2011.0814 |
| *Aedes (Rampamyia) notoscriptus* | Australia | NA | Wilkerson et al. 2021 | https://doi.org/10.1353/book.79680 |
| *Aedes (Stegomyia) aegypti aegypti* | Africa | NA | Powell et al. 2018 | https://doi.org/10.1093/biosci/biy119 |
| *Aedes (Stegomyia) albopictus* | Asia | NA | Medlock et al. 2015 | https://doi.org/10.1017/S0007485315000103 |
| *Aedes (Stegomyia) polynesiensis* | Australia | NA | Wilkerson et al. 2021 | https://doi.org/10.1353/book.79680 |
| *Aedes (Stegomyia) scutellaris* | Asia/Australia | NA | Wilkerson et al. 2021 | https://doi.org/10.1353/book.79680 |
| *Aedes (Stegomyia) unilineatus* | Old World | Africa/Asia | Wilkerson et al. 2021 | https://doi.org/10.1353/book.79680 |
| *Aedes (Tanakaius) togoi* | Asia | NA | Wilkerson et al. 2021 | https://doi.org/10.1353/book.79680 |
| *Anopheles (Anopheles) bancroftii* | Asia/ Australia | NA | Wilkerson et al. 2021; Beebe et al. 2001 | https://doi.org/10.1353/book.79680; https://doi.org/10.1046/j.0962-1075.2001.00278.x |

| | | | | |
|---|---|---|---|---|
| *Anopheles (Anopheles) barbirostris* | Asia | NA | Wilkerson et al. 2021; Taai & Harbach 2015; White 1989 | https://doi.org/10.1353/book.79680; https://doi.org/10.1111/zoj.12236; http://www.ciesin.columbia.edu/docs/001-613/001-613.html |
| *Anopheles (Anopheles) coustani* | Africa | NA | Wilkerson et al. 2021 | https://doi.org/10.1353/book.79680 |
| *Anopheles (Anopheles) crucians* | North America | NA | Wilkerson et al. 2021 | https://doi.org/10.1353/book.79680 |
| *Anopheles (Anopheles) maculipennis* | Europe | NA | Wilkerson et al. 2021; Novikov & Vaulin 2014 | https://doi.org/10.1353/book.79680; http://www.parasitesandvectors.com/content/7/1/389 |
| *Anopheles (Anopheles) punctipennis* | North America | NA | Furumizo et al. 2005; Wilkerson et al. 2021 | http://dx.doi.org/10.2987/8756-971X(2005)21[225:FCOAAP]2.0.CO;2; https://doi.org/10.1353/book.79680 |
| *Anopheles (Anopheles) sacharovi* | Old World | Asia /Europe | White 1989 | http://www.ciesin.columbia.edu/docs/001-613/001-613.html |
| *Anopheles (Cellia) arabiensis* | Africa | NA | Wilkerson et al. 2021; White 1989 | https://doi.org/10.1353/book.79680; http://www.ciesin.columbia.edu/docs/001-613/001-613.html |
| *Anopheles (Cellia) funestus* | Africa | NA | Wilkerson et al. 2021, White 1989 | https://doi.org/10.1353/book.79680; http://www.ciesin.columbia.edu/docs/001-613/001-613.html |
| *Anopheles (Cellia) gambiae s.l.* | Africa | NA | Wilkerson et al. 2021, White 1989 | https://doi.org/10.1353/book.79680; http://www.ciesin.columbia.edu/docs/001-613/001-613.html |
| *Anopheles (Cellia) litoralis* | Asia | NA | Wilkerson et al. 2021 | https://doi.org/10.1353/book.79680 |
| *Anopheles (Cellia) pharoensis* | Africa | NA | Wilkerson et al. 2021, White 1989 | https://doi.org/10.1353/book.79680; http://www.ciesin.columbia.edu/docs/001-613/001-613.html |
| *Anopheles (Cellia) stephensi* | Asia | NA | Faulde et al. 2014; Sinka et al. 2020 | https://doi.org/10.1016/j.actatropica.2014.06.016; https://www.pnas.org/doi/abs/10.1073/pnas.2003976117 |
| *Anopheles (Cellia) subpictus* | Asia | NA | Chandra et al. 2010 | https://doi.org/10.1016/j.actatropica.2010.02.005 |
| *Anopheles (Nyssorhynchus) albimanus* | Americas | NA | Wilkerson et al. 2021 | https://doi.org/10.1353/book.79680 |
| *Anopheles (Nyssorhynchus) darlingi* | Americas | NA | White 1989; Mirabello & Conn 2006 | http://www.ciesin.columbia.edu/docs/001-613/001-613.html; https://www.nature.com/articles/6800805.pdf |
| *Armigeres (Armigeres) subalbatus* | Asia | NA | Wilkerson et al. 2021 | https://doi.org/10.1353/book.79680 |
| *Culex (Barraudius) modestus* | Old World | Africa/Asia / Europe | Soto & Delang 2023 | https://doi.org/10.1186/s13071-023-05997-6 |
| *Culex (Culex) annulirostris* | Asia/ Australia | NA | Wilkerson et al. 2021 | https://doi.org/10.1353/book.79680 |
| *Culex (Culex) antennatus* | Asia | NA | Wilkerson et al. 2021 | https://doi.org/10.1353/book.79680 |
| *Culex (Culex) coronator* | Americas | NA | Wilkerson et al. 2021; Romero-Weaver 2023 | https://doi.org/10.1353/book.79680; https://doi.org/10.32473/edis-IN1385-2022 |
| *Culex (Culex) fuscocephala* | Asia | NA | Wilkerson et al. 2021 | https://doi.org/10.1353/book.79680 |

| | | | | |
|---|---|---|---|---|
| *Culex (Culex) gelidus* | Asia | NA | Wilkerson et al. 2021; Sudeep 2014 | https://doi.org/10.1353/book.79680; https://pubmed.ncbi.nlm.nih.gov/25540955/ |
| *Culex (Culex) pipiens s.l.* | Africa | NA | Farajollahi et al. 2011 | https://doi.org/10.1016/j.meegid.2011.08.013 |
| *Culex (Culex) quinquefasciatus* | Asia | NA | Farajollahi et al. 2011; Fonesca et al. 2006 | https://doi.org/10.1016/j.meegid.2011.08.013; https://doi.org/10.4269/ajtmh.2006.74.284 |
| *Culex (Culex) sitiens* | unknown | Africa/Asia-Trop/ Australia | Wilkerson et al. 2021 | https://doi.org/10.1353/book.79680 |
| *Culex (Culex) tarsalis* | North America | NA | Wilkerson et al. 2021 | https://doi.org/10.1353/book.79680 |
| *Culex (Culex) tritaeniorhynchus* | Old World | Africa/Asia | Alves et al. 2014; Wilkerson et al. 2021 | http://www.scvz.org/zoolcv/vol5no1/Alves%20et%20al%20Culex%20tritaeniorhynchus.pdf; https://doi.org/10.1353/book.79680 |
| *Culex (Oculeomyia) bitaeniorhynchus* | unknown | Africa/Asia Australia | Sirivanakarn 1973 | https://apps.dtic.mil/sti/pdfs/ADA511697.pdf |
| *Mansonia (Mansonia) titillans* | Americas | NA | Wilkerson et al. 2021 | https://doi.org/10.1353/book.79680 |
| *Mansonia (Mansonioides) uniformis* | unknown | Africa/Asia-Trop/ Australia | White 1989 | http://www.ciesin.columbia.edu/docs/001-613/001-613.html |

**Suppl. Material Tab. 6.** Species names, descriptors, abbreviations used in the manuscript, and whether the species was introduced or also established outside its native region.

| Species | Descriptor | Abbreviation | Established outside their native range |
|---|---|---|---|
| *Aedes (Aedimorphus) vexans* | Theobald, 1903 | *Ae. vexa* | yes |
| *Aedes (Fredwardsius) vittatus* | Bigot, 1861 | *Ae. vitt* | yes |
| *Aedes (Georgecraigius) atropalpus* | Coquillett, 1902 | *Ae. atro* | yes |
| *Aedes (Hulecoeteomyia) japonicus japonicus* | Theobald, 1901 | *Ae. japo* | yes |
| *Aedes (Ochlerotatus) camptorhynchus* | Thomson, 1869 | *Ae. camp* | no |
| *Aedes (Ochlerotatus) mcintoshi* | Huang, 1985 | *Ae. mcin* | no |
| *Aedes (Ochlerotatus) vigilax* | Skuse, 1889 | *Ae. vigi* | yes |
| *Aedes (Protomacleaya) triseriatus* | Say, 1823 | *Ae. tris* | no |
| *Aedes (Rampamyia) notoscriptus* | Skuse, 1889 | *Ae. noto* | yes |
| *Aedes (Stegomyia) aegypti aegypti* | Linnaeus, 1762 | *Ae. aegy* | yes |
| *Aedes (Stegomyia) albopictus* | Skuse, 1895 | *Ae. albo* | yes |
| *Aedes (Stegomyia) polynesiensis* | Marks, 1951 | *Ae. poly* | no |
| *Aedes (Stegomyia) scutellaris* | Walker, 1859 | *Ae. scut* | no |
| *Aedes (Stegomyia) unilineatus* | Theobald, 1906 | *Ae. unil* | yes |
| *Aedes (Tanakaius) togoi* | Theobald, 1907 | *Ae. togo* | yes |
| *Anopheles (Anopheles) bancroftii* | Giles, 1902 | *An. banc* | yes |
| *Anopheles (Anopheles) barbirostris* | van der Wulp, 1884 | *An. barb* | no |
| *Anopheles (Anopheles) coustani* | Laveran, 1900 | *An. cous* | no0 |
| *Anopheles (Anopheles) crucians* | Wiedemann, 1828 | *An. cruc* | no |
| *Anopheles (Anopheles) maculipennis* | Meigen, 1818 | *An. macu* | no |
| *Anopheles (Anopheles) punctipennis* | Say, 1823 | *An. punc* | no |
| *Anopheles (Cellia) arabiensis* | Patton, 1905 | *An. arab* | no |
| *Anopheles (Cellia) funestus* | Giles, 1900 | *An. fune* | no |
| *Anopheles (Cellia) gambiae s.l.* | Giles, 1900 | *An. gamb* | yes |
| *Anopheles (Cellia) litoralis* | King, 1932 | *An. lito* | no |
| *Anopheles (Cellia) pharoensis* | Theobald, 1901 | *An. phar* | no |
| *Anopheles (Cellia) stephensi* | Liston, 1901 | *An. step* | yes |
| *Anopheles (Cellia) subpictus* | Grassi, 1899 | *An. subp* | yes |
| *Anopheles (Nyssorhynchus) albimanus* | Wiedemann, 1821 | *An. albi* | yes |
| *Anopheles (Nyssorhynchus) darlingi* | Root, 1926 | *An. darl* | yes |
| *Armigeres (Armigeres) subalbatus* | Coquillett, 1898 | *Ar. suba* | yes |

| | | | |
|---|---|---|---|
| *Culex (Barraudius) modestus* | Ficalbi, 1890 | *Cx. mode* | yes |
| *Culex (Culex) annulirostris* | Skuse, 1889 | *Cx. annu* | no |
| *Culex (Culex) antennatus* | Becker, 1903 | *Cx. ante* | no |
| *Culex (Culex) coronator* | Dyar & Knab, 1906 | *Cx. coro* | yes |
| *Culex (Culex) fuscocephala* | Theobald, 1907 | *Cx. fusc* | yes |
| *Culex (Culex) gelidus* | Theobald, 1901 | *Cx. geli* | yes |
| *Culex (Culex) pipiens s.l.* | Linnaeus, 1758 | *Cx. pipi* | yes |
| *Culex (Culex) quinquefasciatus* | Say, 1823 | *Cx. quin* | yes |
| *Culex (Culex) sitiens* | Wiedemann, 1828 | *Cx. siti* | yes |
| *Culex (Culex) tarsalis* | Coquillett, 1896 | *Cx. tars* | yes |
| *Culex (Culex) tritaeniorhynchus* | Giles, 1901 | *Cx. trit* | yes |
| *Culex (Oculeomyia) bitaeniorhynchus* | Giles, 1901 | *Cx. bita* | no |
| *Mansonia (Mansonia) titillans* | Walker, 1848 | *Ma. titi* | no |
| *Mansonia (Mansonioides) uniformis* | Theobald, 1901 | *Ma. unif* | yes |

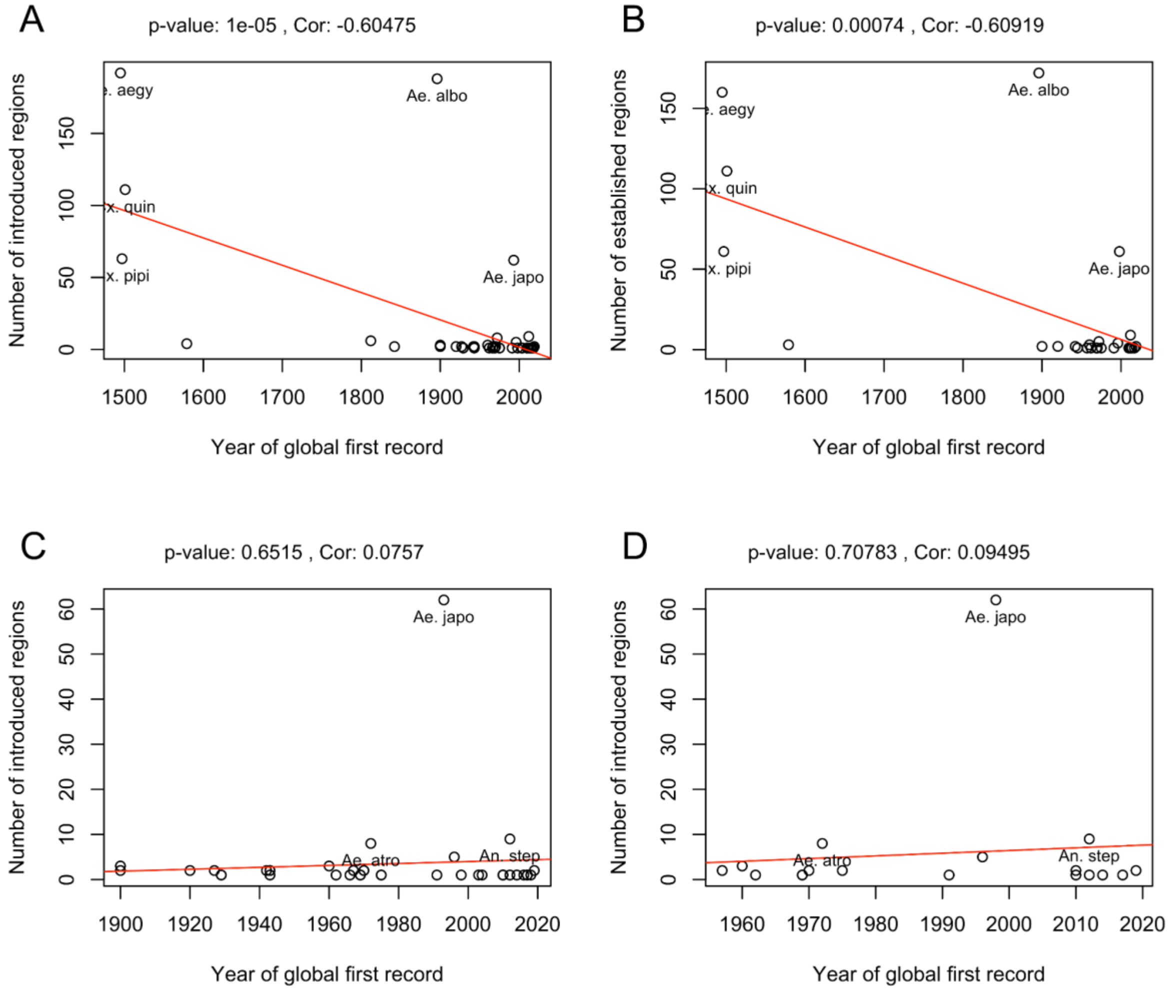


**Suppl. Material Fig. 5.** Pearson correlation test between the timing of spread and number of invaded regions for all introduced species (A), all introduced established species (B), all introduced species with global first record after 1900 (C) and after 1950 (D).

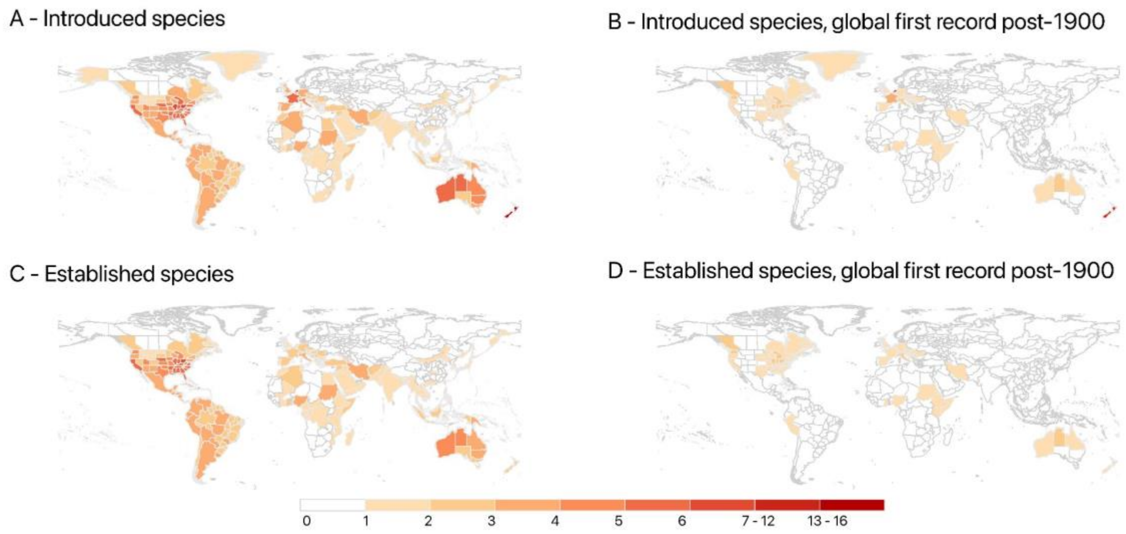


**Suppl. Material Fig. 6.** Regional level richness of introduced non-native species. Maps represent introduced species richness (A), introduced species with first global record post-1900 (B), established species richness (C), and established species with first global record post-1900 (D).

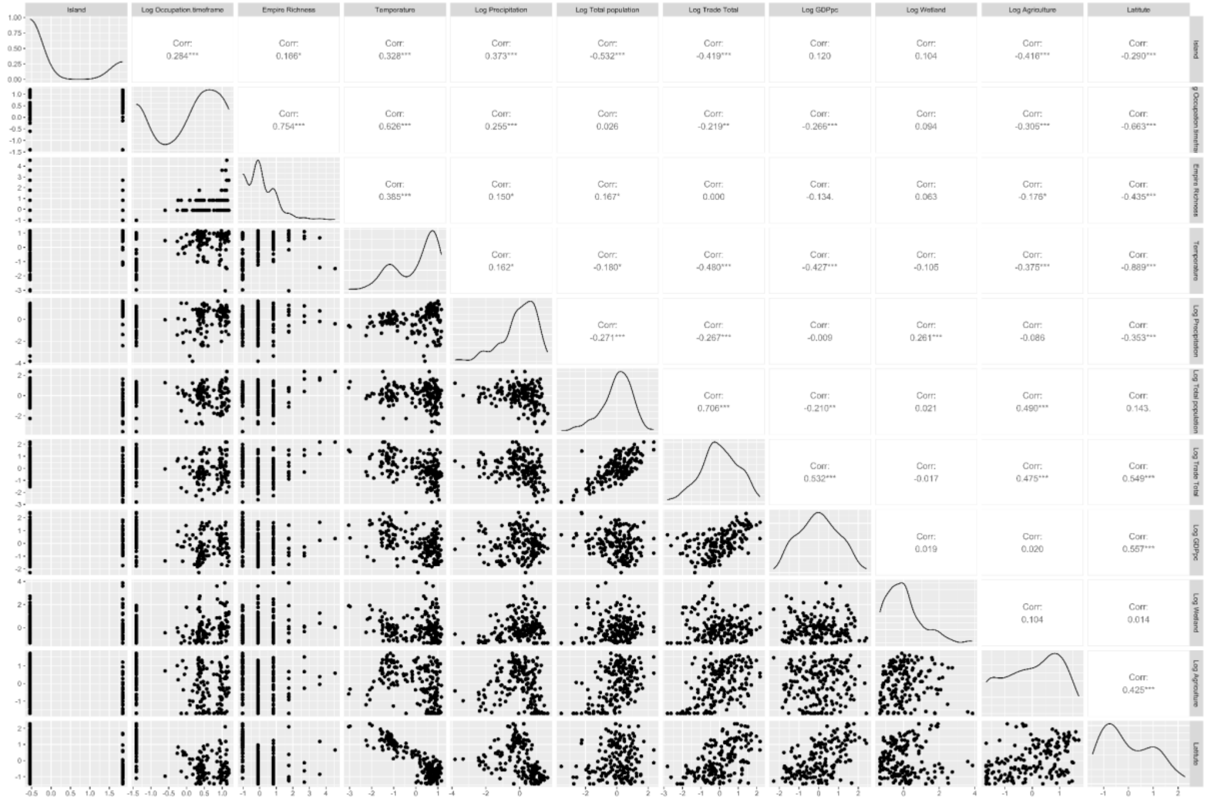


**Suppl. Material Fig. 7.** Correlation matrix for the predictors used in the models to test the drivers. Variable description:Log Occupation.timeframe: Time a country was colonized by European colonial powers log transformed; Empire.Richness: Number of colonial empires that occupied the country; Temperature: average yearly temperature between 1970 and 2000 in °C; Log GDPpc: average GDP per capita between 2017 and 2021; Log Precipitation: yearly average precipitation between 1970 and 2000 in mm log transformed; Log Total population: average total population between 2017 and 2021 log transformed; Log Trade Total: average between import and export of trade as total US$ between 2017 and 2021 log transformed; Log Wetland: percentage of the country's area covered by wetlands log transformed; Log Agriculture: percentage of the country's area covered by agriculture log transformed; Latitude: absolute latitude indicating only the magnitude of the distance from the Equator without specifying the direction (north or south)
